# Peculiar features of the plastids of the colourless alga *Euglena longa* and photosynthetic euglenophytes unveiled by transcriptome analyses

**DOI:** 10.1101/358895

**Authors:** Kristína Záhonová, Zoltán Füssy, Erik Birčák, Anna M. G. Novák Vanclová, Vladimír Klimeš, Matej Vesteg, Juraj Krajčovič, Miroslav Oborník, Marek Eliáš

## Abstract

**Background:** Euglenophytes are an interesting algal group that emerged within the ancestrally plastid-lacking Euglenozoa phylum by acquiring a plastid from a green algal donor. However, the knowledge of euglenophyte plastid biology and evolution is highly incomplete, partly because euglenophytes have so far been little studied on a genome- and transcriptome-wide scale. Transcriptome data from only a single species, *Euglena gracilis*, have been exploited to functional insights, but aspects of the plastid biology have been largely neglected.

**Results:** To expand the resources for studying euglenophyte biology and to improve our knowledge of the euglenophyte plastid function and evolution, we sequenced and analysed the transcriptome of the non-photosynthetic species *Euglena longa*. The transcriptomic data confirmed the absence of genes for the photosynthetic machinery in this species, but provided a number of candidate plastid-localized proteins bearing the same type of N-terminal bipartite topogenic signals (BTSs) as known from the photosynthetic species *E. gracilis*. Further comparative analyses using transcriptome assemblies available for *E. gracilis* and two additional photosynthetic euglenophytes of the genus *Eutreptiella* enabled us to unveil several salient aspects of the basic plastid infrastructure in euglenophytes. First, a number of plastidial proteins seem to reach the organelle as C-terminal translational fusions with other BTS-bearing proteins. Second, the conventional eubacteria-derived plastidial ribosomal protein L24 is missing and seems to have been replaced by very different homologs of the archaeo-eukaryotic origin. Third, no homologs of any key component of the TOC/TIC system (translocon of the outer/inner chloroplast membrane) and the plastid division apparatus are discernible in euglenophytes, and the machinery for intraplastidial protein targeting has been simplified by the loss of the cpSRP/cpFtsY system and the SEC2 translocon. Lastly, euglenophytes proved to encode a plastid-targeted homolog of the termination factor Rho horizontally acquired from a Lambdaproteobacteria-related donor, suggesting an unprecedented modification of the transcription mechanism in their plastid.

**Conclusions:** Our study suggests that the euglenophyte plastid has been substantially remodelled comparted to its green algal progenitor by both loss of original and acquisition of novel molecular components, making it a particularly interesting subject for further studies.

## Background

Euglenophytes, exemplified by the highly studied mixotrophic alga *Euglena gracilis*, are a peculiar algal group constituting one of the many lineages of the phylum Euglenozoa [1]. Euglenophytes and other euglenozoans share many unusual features, such as regulation of gene primarily at the post-transcriptional level [2–4]. Furthermore, euglenozoans employ *trans*-splicing to process mRNA molecules, whereby the 5’-end of pre-mRNA is replaced by the 5’-end of the specialized spliced leader (SL) RNA, resulting in the presence of the invariant SL sequence at the 5’-end of mature mRNAs [5, 6].

Despite their interesting biology, euglenophytes have not yet been properly studied by genome-wide approaches. A few studies employed transcriptome sequencing of *E. gracilis* to investigate particular aspects of its gene repertoire and selected functional pathways [7–9]. Two independent transcriptome assemblies are available in the GenBank database, and a nuclear genome draft has been announced, although not yet made public at the time of writing of this paper [7]. In addition, transcriptome assemblies of two different isolates of the marine euglenophyte genus *Eutreptiella* (*E. gymnastica* NIES-381 and *E. gymnastica*-like CCMP1597) were sequenced as part of the MMETSP project [10], but no specific analyses of this data resource have been reported. Hence, further studies are clearly needed to improve our understanding of the molecular underpinnings of the euglenophyte life and evolution.

The defining feature of euglenophytes is a complex three-membrane-bounded plastid derived from a green alga belonging to Pyramimonadales [1, 11–13]. As in other plastid-bearing eukaryotes, only a minority of plastid proteins are encoded by the plastid genome; the nucleus-encoded majority then need to cross the three membranes of the euglenophyte plastid envelope to reach the site of their function. The mechanism of protein targeting to the plastid has been partially characterized in *E. gracilis* [14]. The proteins co-translationally enter the endoplasmic reticulum (ER) and are transported further by vesicular trafficking, passing the Golgi apparatus *en route* to the plastid. The *E. gracilis* plastid-targeted proteins bear a discernible presequence, an N-terminal bipartite topogenic signal (BTS), which comes in two main variants [15]. Both include an N-terminal signal peptide mediating the import into the ER, followed by a plastid transit peptide that is exposed upon signal peptide cleavage and mediates import across the two inner chloroplast membranes. In Class I presequences, the transit peptide is followed by a transmembrane domain (TMD) that anchors the transported protein in the membrane during its subcellular relocation, whereas Class II presequences lack the anchoring TMD. The signal peptide itself typically has physicochemical properties of a TMD, resulting in a characteristic double-TMD motif in the Class I presequences of euglenophyte plastid proteins [15]. The characteristic structure of the *E. gracilis* plastid-targeting BTSs has facilitated *in silico* identification of candidates for plastid-targeted proteins in euglenophytes [15–18]. However, the proteome of neither euglenophyte plastid has yet been reconstructed in full.

Although most euglenophytes are photosynthetic, several lineages independently lost photosynthesis and became secondarily heterotrophic [19]. The fate of their plastid is generally unknown, except for *Euglena longa* (originally described as *Astasia longa*), where a nonphotosynthetic plastid has been preserved, as evident from the presence of a plastid genome sequenced a long time ago [20]. We have recently demonstrated that an intact plastid genome is essential for the *E. longa* survival, in contrast to photosynthetic *E. gracilis* [21]. The *E. longa* plastid genome size (75 kbp) is approximately half of that of *E. gracilis*, with the difference attributed primarily to the absence of photosynthesis-related genes. The only exception is the *rbcL* gene encoding the large subunit of the enzyme ribulose-1,5-bisphosphate carboxylase/oxygenase (RuBisCO LSU) retained in the *E. longa* plastid genome [20, 22]. However, the plastid organelle itself remains elusive. Double-membrane bodies similar to those present in dark-grown *E. gracilis* were observed in *E. longa* and interpreted as plastids [23], but this identification is uncertain given that euglenophyte plastids studied in detail possess three bounding membranes [1]. Likewise, the physiological role of the *E. longa* plastid remains unknown.

It is conceivable that a lot could be learned about the *E. longa* plastid by analysing its genome sequence. However, the genome sequencing project initiated by others for *E. gracilis* revealed that the genome of this species is huge and difficult to assemble [7]. As a close relative [19, 24], *E. longa* most likely shares general genomic features with *E. gracilis*, making genome sequencing impractical. Hence, to build a resource for exploration of the plastid function and other aspects of the *E. longa* biology, we sequenced and assembled the transcriptome of this species. The sequencing data were obtained from cultures grown at two different light regimes (in the dark and in the light) to improve the coverage of differentially expressed genes and to maximize the chance we detect possible traces of genes encoding the photosynthesis-related machinery. Data extracted from the sequenced *E. longa* transcriptome proved instrumental in characterizing the function of the RuBisCO enzyme in this species [22] and enabled identification of a novel Euglenozoa-specific form of the Rheb GTPase [25], but publication of the whole transcriptome assembly has been pending.

Here we describe the general characteristics of the transcriptome and demonstrate its utility for unravelling the biology of the *E. longa* plastid. We provide the first insights into the basic infrastructure of the plastid and compare it to the molecular machinery of plastid biogenesis in photosynthetic euglenophytes. Our results not only *E. longa*-specific simplification related to the loss of photosynthesis, but also a surprising reduction of the plastid biogenesis machinery in euglenophytes in general. Finally, we report on a case of an expansion of the euglenophyte plastid functions by acquisition of a plastid-targeted homolog of the bacterial transcription termination factor Rho, which is an unprecedented feature among all plastid-bearing eukaryotes studied to date. We believe that the *E. longa* transcriptome, now made available to the whole scientific community, will become an important resource for further research of various aspects of euglenophyte biology.

## Results and Discussion

### The transcriptome of *Euglena longa*: not all mRNAs bear the 5’ end *trans*-spliced leader sequence

Our transcriptomic assembly of *E. longa* resulted in 65,563 transcript models, a number somewhat smaller than the numbers reported in recent transcriptomic studies for *E. gracilis* (113,152 [9]; 72,506 [7]). Since different sequence assembly algorithms employed certainly contribute to the difference in contig numbers, we carried out a BUSCO search for conserved unique eukaryote orthologs to assess the quality of our data. 89.1% of BUSCO genes were found to be complete in our dataset, and further 4.3% of orthologs to be present as fragments. These characteristics are similar to those of *E. gracilis* transcriptomic data (Additional file 1: Figure S1; [7, 9]) and suggest that our assembly covers the majority of genes in the *E. longa* genome.

31,742 *E. longa* transcript models (46%) possessed at least a partial spliced leader (SL) sequence as a sign of their 5’-end completeness. The percentage of transcripts with the SL sequence in *E. gracilis* was comparable (54%; [9]), even though SL absence was so far directly experimentally demonstrated only for mRNA of a single *E. gracilis* gene, the one encoding the nucleolar protein fibrillarin [26]. Since *in silico* prediction of protein subcellular localisation requires full-length protein sequences, it was critical to understand whether the high proportion of SL sequence-lacking transcripts implies a high fraction of truncated sequences. Therefore, we chose candidate transcripts that lack the SL sequence in our transcriptome assembly (listed in Additional file 2: Table S1) and tested the presence of the SL sequence at the 5’-end of the respective mRNAs using PCR (with cDNA as the template). Several transcripts failed to be amplified when using the SL-specific forward primer (Fig. 1A, lane S), but were amplified when gene-specific forward primers were used (Fig. 1A, lane X), indicating that they truly lack the SL sequence.

**Fig. 1:**
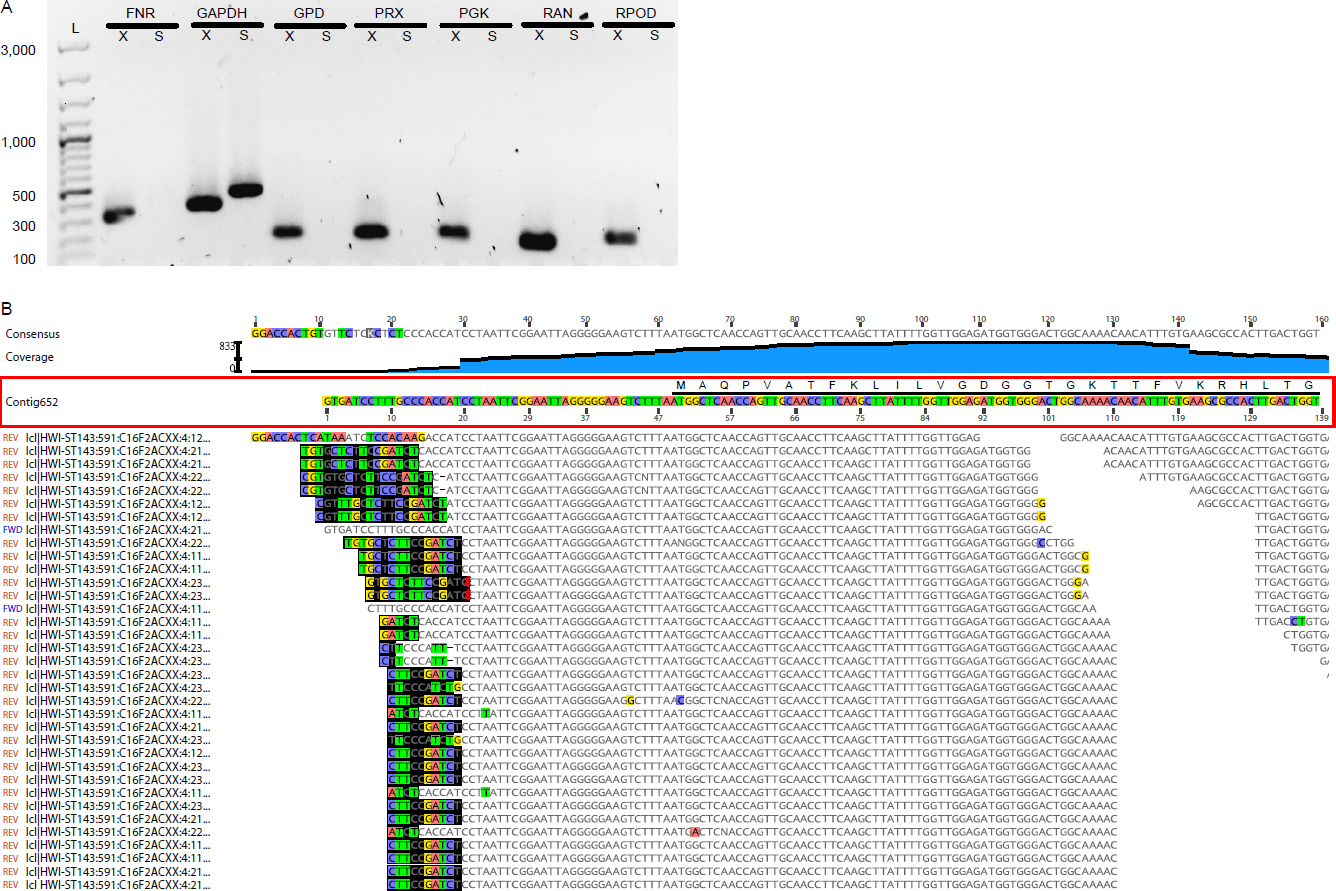
Presence/absence of the spliced leader (SL) in selected transcripts in *E. longa*. A) PCR (with cDNA as the template) with an SL-specific forward primer was done to assay whether the lack of SL in exemplar assembled contigs (listed in Additional file 2: Table S2) mirrors real SL absence in the respective transcripts. Lanes X show PCR products with gene-specific primers (positive control), lanes S show products with SL forward and gene-specific reverse primers. Lane L is a 100-bp size ladder with sizes shown for selected bands. Assayed transcripts: FNR, ferredoxin-NADP+ reductase; GAPDH, glyceraldehyde-3-phosphate dehydrogenase (positive control for the SL-specific primer); GPD, glycerol-phosphate dehydrogenase; PRX, peroxiredoxin; PGK, phosphoglycerate kinase; RAN, RAN GTPase; RPOD, RNA polymerase sigma factor. B) Mapping of raw sequence reads to the 5’-end of the RAN GTPase transcript from *E. longa* confirms the absence of the SL sequence. Untrimmed Illumina primer sequences at the 5’-end of several reads are highlighted by black background. The coding sequence is shown by the black horizontal line and the amino acid sequence above the Contig652 nucleotide sequence. The apparent discontinuity in the read coverage (around the position 110 of the contig) is not real and stems from omission from the figure of many read mapping to this region (to make the scheme smaller).

To further test the notion that some *E. longa* mRNAs may not undergo *trans*-splicing, we chose the highly expressed and conserved gene for the GTPase RAN (implicated in nucleocytoplasmic transport; [27]) and attempted to extend the SL sequence-lacking 5’-end of the respective transcript contig by using all available RNA-seq reads. While the read coverage of the 5’-end is high, no extension to recruit the SL sequence to the end is possible (Fig. 1B). Hence, the maturation of the mRNA 5’-end by SL *trans*-splicing is not universal to all genes in euglenophytes. However, not all SL-sequence lacking contigs in the transcriptome assembly necessarily attest to the lack of *trans*-splicing, as some of them are truly truncated and others may represent un-spliced variants of normally *trans*-spliced mRNAs. Indeed, we found examples of both cases (Additional file 1: Figure S2). The genome sequence of *E. gracilis* that should soon become available will enable to carry out a systematic analysis of the occurrence of SL *trans-*splicing across the transcriptome.

### No traces of the photosynthetic machinery in the transcriptome of *Euglena longa*

Although polyA-selection was employed during the RNA-seq library preparation, we detected in our assembly transcripts of some genes of the plastid genome (Additional file 1: Figure S3; Additional file 2: Table S2). This may mean that some plastid mRNAs are polyadenylated in *E. longa*, as demonstrated for *E. gracilis* [28], but inefficient removal of non-polyadenylated RNA molecules cannot be ruled out as an alternative explanation. Comparison of the transcript sequences with the plastid genome sequence [20] revealed that the second intron in the *rps2* gene has an incorrectly delimited 3’-border in the current genome annotation, rendering the coding sequence shorter by one amino acid. No signs of plastid mRNA editing were observed.

The *E. longa* plastid genome lacks many of the genes found in plastid genomes of other euglenophytes (Additional file 1: Figure S3). Except for the *rps18* gene encoding the ribosomal protein S18, all missing genes code for components of the photosynthetic machinery, i.e. photosystems I and II, the cytochrome *b*_6_*f* complex, membrane ATP synthase, and the enzyme Mg-protoporphyrin IX chelatase involved in chlorophyll synthesis. We searched the *E. longa* transcriptome assembly to investigate possible transfer of these genes into the nuclear genome, but we did not find any of them, suggesting that no endosymbiotic gene transfer occurred in the *E. longa* lineage after its separation from the *E. gracilis* lineage. We likewise failed to find homologs of other conserved components of the main photosynthetic complexes encoded by the nuclear genome in *E. gracilis* or other photosynthetic euglenophytes (Additional file 2: Table S2). We assume that if the photosynthesis-related genes were present in the *E. longa* nuclear genome, we would detect transcripts of at least some of them, since one of the sequenced cultures was grown in the light. This corroborates that photosynthesis is truly missing in *E. longa*.

The apparent loss of the gene for the plastid ribosomal protein S18, otherwise broadly conserved in photosynthetic eukaryotes [29], raises the question how the plastid ribosome small subunit is affected by the absence of this protein. However, *rps18* is missing from some other colourless plastid genomes, such as those of apicomplexans, the chlorophytes *Helicosporidium* sp., *Prototheca stagnora*, and *Polytoma uvella*, and the diatom *Nitzschia* sp. NIES-3581 [30–32]. Apicomplexans were reported to lack even a nucleus-encoded apicoplast-targeted version of S18 (while keeping a mitochondrion-targeted version; [33]), and we likewise could not find a plastid-targeted S18 protein in the available nuclear genome data from *Helicosporidium*, suggesting that S18 is not absolutely essential for translation in the plastid.

### Probing for nucleus-encoded plastid proteins in *E. longa*: aminoacyl-tRNA synthetases and ribosomal proteins

We previously reported several *E. longa* proteins presumed to be targeted to its plastid, including the small RuBisCO subunit (RBCS), RuBisCO activase, the chaperonins GroEL/GroES, and the assembly factor RAF ([22]; see also Additional file 2: Table S3), but we did not analyse their targeting sequences in detail. To get a broader representative set of nucleus-encoded proteins likely imported into the *E. longa* plastid, we searched the transcriptome assembly for proteins from two functional categories expected to be present in the *E. longa* plastid: (1) aminoacyl-tRNA synthetases needed for charging tRNAs specified by the plastid genome; and (2) ribosomal proteins not encoded by the plastid genome. The same search was done for *E. gracilis* to further assess the representativeness of the *E. longa* transcriptome assembly with special regard to nuclear genes encoding plastidial proteins. Plastidial aminoacyl-tRNA synthetases and ribosomal proteins were discriminated from homologs functioning in other compartments (the cytosol and the mitochondrion) by virtue of their closer sequence similarity to plastidial homologs in other eukaryotes and/or by the presence of an N-terminal extension bearing general characteristics of BTSs as previously defined in *E. gracilis*.

We found the same set of these two categories of proteins in both species, although in several cases the respective sequence was 5’-truncated in one or the other species (Additional file 2: Table S4 and S5). Some of these incomplete sequences could be extended by manual iterative recruitment of sequencing reads to the 5’-end of the transcript (often up to the SL sequence), providing the missing part of the coding sequence and consequently the BTS in the encoded protein. Although sequences of some putative plastid-targeted proteins remained truncated even when using this approach, their plastid localisation can be assumed based on the premise that the missing N-terminal region of the protein has the features as its ortholog from the other *Euglena* species. With this premise, plastidial aminoacyl-tRNA synthetases cognate to all twenty amino acids exist in both *Euglena* species, two of them being included in one fusion protein (see below). The presence of a plastidial version of Gln-tRNA synthetase in *Euglena* spp. is noteworthy, because plastids of different plant and algal groups lack this enzyme and instead rely on an alternative two-step process of Gln-tRNA synthesis inherited from the cyanobacterial progenitor of the plastid and mediated by Glu-tRNA synthetase and Glu-tRNA amidotransferase [34, 35]. However, putative plastid-targeted glutaminyl-tRNA synthetases were recently found in diatoms and the cryptophyte *Guillardia theta* [36], so plastids may be more diverse in their mechanism of Gln-tRNA synthesis than previously thought. The euglenophyte plastid-targeted Gln-tRNA synthetase was apparently gained by HGT from a bacterium, most likely a member of Gammaproteobacteria, judging from the identity of the best blastp hits (data not shown).

An even more interesting picture emerged from the analysis of plastidial ribosomal proteins. The set of ribosomal proteins with apparent plastid-targeting presequences identified in the transcriptome data complemented the set encoded by the plastid genomes, such that all subunits of the plastid ribosome known from other algal groups (see [33]) are conserved in both *Euglena* species, with two exceptions. The first is the lack of S18 in *E. longa* discussed in the previous section, and the second is the absence of the expected eubacteria-like L24. Both *Euglena* species each instead encode two other proteins of the same ortholog family (called uL24 according to the latest unified nomenclature of ribosomal proteins; [37]) that exhibit N-terminal extensions fitting the structure of the BTS (Additional file 2: Table S5). The two *Eutreptiellla* species each possess only one such protein, which however represent two different paralogs that originated before the euglenophyte radiation (with possible subsequent differential loss in the *Eutreptiella* lineage; Fig. 2). These two paralogs share a common ancestor apparently belonging to the archaeo-eukaryotic uL24 family branch, often called L26. A phylogenetic analysis did not suggest a more specific scenario, as the position of the paralog pair outside the eukaryotic and archaeal clades in the tree (Fig. 2) is most likely due to their rapid divergence erasing the phylogenetic signal in these short proteins.

**Fig. 2:**
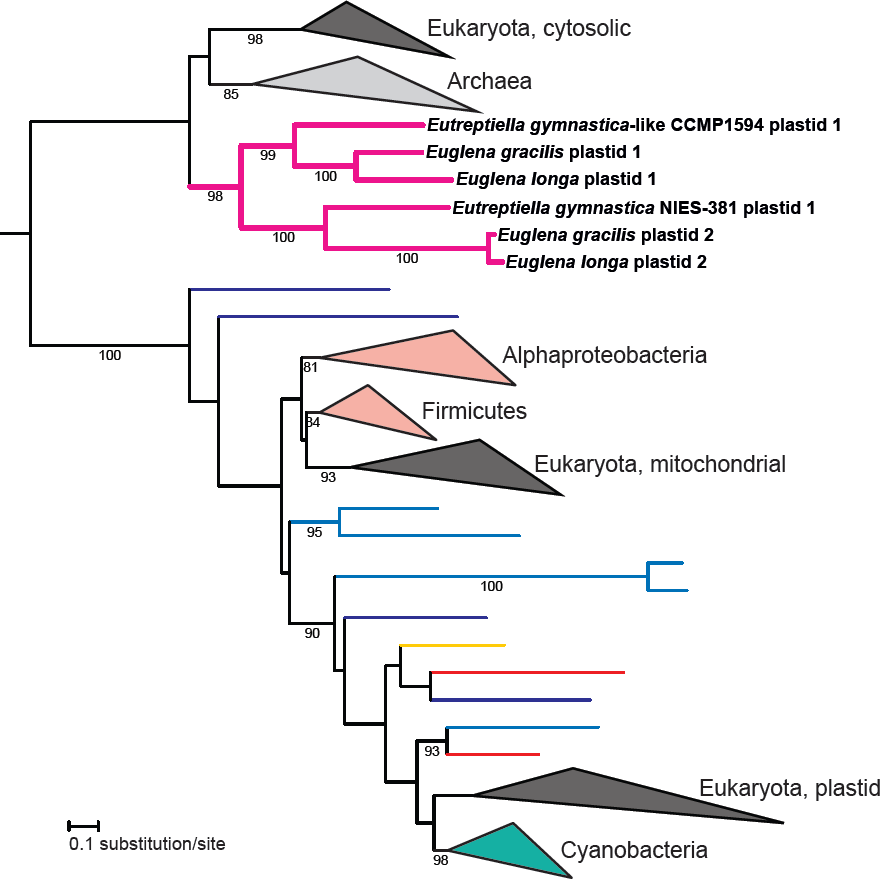
**Phylogenetic analysis of the uL24 family of ribosomal proteins.** The tree shows the phylogenetic position of the presumably plastid-localized L26-related proteins in euglenophytes. Bootstrap support values are given when ≥ 80.

It is tempting to speculate that the plastid-targeted L26-related proteins functionally compensate for the absence of the eubacteria-like L24 in the euglenophyte plastidial ribosome, despite considerable sequence divergence between the eubacterial and archaeo-eukaryotic homologs. Such a replacement of an organellar ribosomal protein by a homolog from a different phylogenetic domain may seem unlikely, but is apparently possible. At least two similar cases have been documented, both featuring a novel paralog of the eukaryote-type cytosolic ribosomal protein replacing the homologous eubacteria-like counterpart of the organellar ribosome: the eubacteria-type S8 protein in angiosperm mitochondria replaced by the eukaryotic protein S15A [38] and the ancestral eubacterial L23 replaced by the eukaryotic homolog in the plastid of spinach (but perhaps also in other plants; [39, 40]). The euglenophyte plastidial L26-related proteins thus may be another such case. Why *Euglena* spp. exhibit two different plastid-targeted L26-related proteins is unclear, but it is possible that one paralog is not a part of the ribosome itself and performs another role. Indeed, the cytosolic L26 has a ribosome-independent function as a regulator of translation of p53 family proteins in mammals [41, 42].

### Protein import into the euglenophyte plastids: targeting sequences and translational fusions

The collection of the high-confidence candidates for *E. longa* proteins targeted to the plastid established in our previous study [22] and by the analyses described above enabled us to evaluate the general characteristics of the plastid-targeting presequences in this species. The analysis revealed that the N-terminal extensions of these proteins exhibit the same characteristics as the BTS defined in *E. gracilis* (see above). Specifically, we could find presequences of both class I and class II (Additional file 1: Figure S4), with the relative abundance of class I being higher than in *E. gracilis* (Additional file 2: Table S3 and S4). The class I presequences typically exhibited the pattern of two predicted TMDs separated by a hydrophilic amino acid stretch according to the 60±8 rule [15], and the N-terminal signal peptide was predicted in most proteins by all tools employed. This suggests that both *Euglena* species share a similar route of plastid protein import and, presumably, that the *E. longa* plastid envelope also consists of three membranes.

We previously documented that the RBCS transcript in *E. longa* encodes a polyprotein comprising a single targeting presequence followed by several monomers of the mature RBCS separated by a conserved decapeptide linker [22]. Similar organization is found also in the *E. gracilis* RBCS and is characteristic of a handful of other photosynthesis-related proteins in *E. gracilis* [43–46] and the dinoflagellate *Prorocentrum minimum* [47]. In *E. gracilis*, individual RBCS units are processed by proteolytic cleavage upon import of the polyprotein into the plastid [44], while RBCS polymer processing was not detected in *E. longa [22]*. Interestingly, we now observed that the translation elongation factor EF-Ts is encoded in a similar fashion in both *E. longa* and *E. gracilis*, with a plastid-targeting presequence followed by two EF-Ts monomers separated by a linker (Fig. 3).

**Fig. 3:**
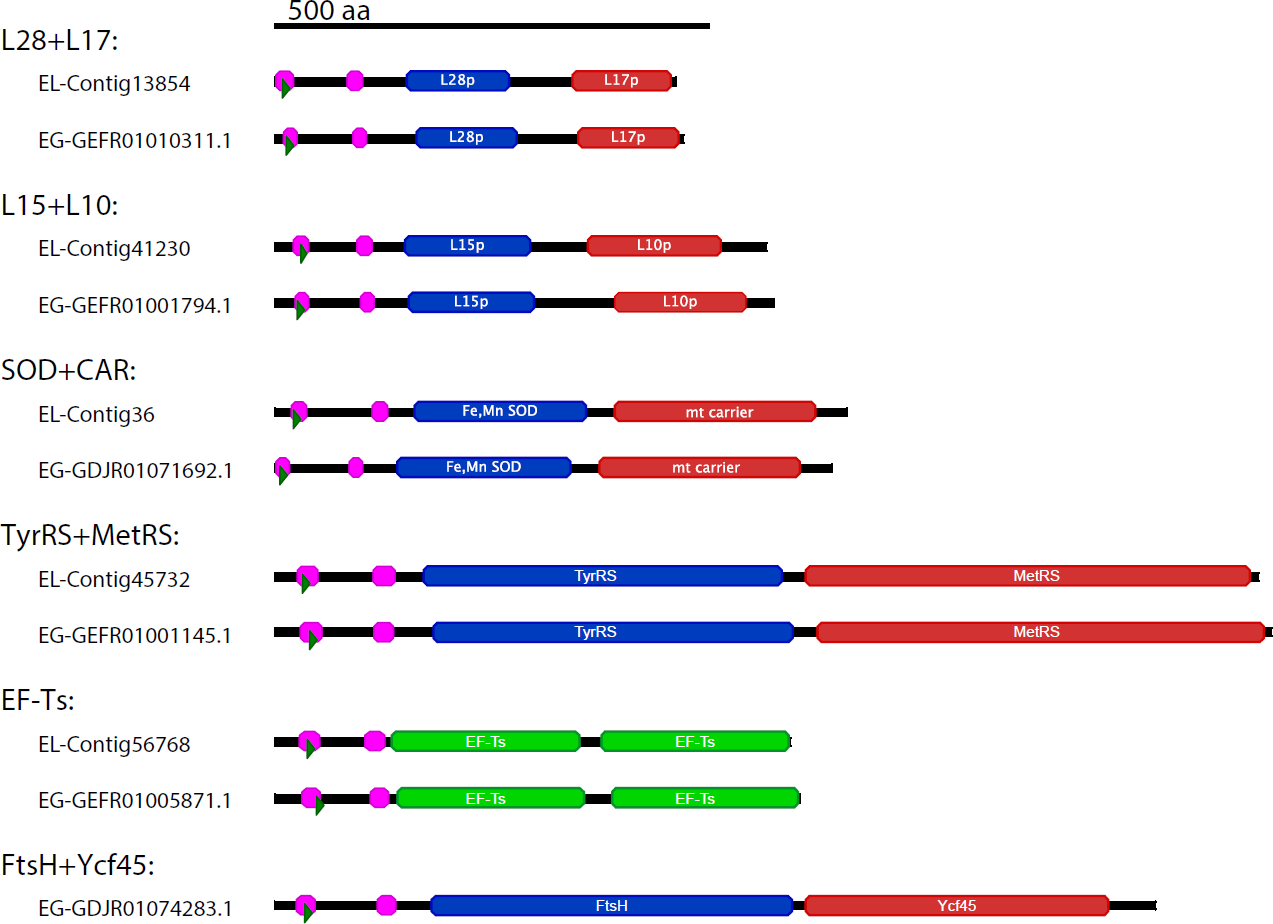
Domain structure of fusion proteins in *E. longa* and *E. gracilis*. Regions corresponding to separate mature proteins presumably released by processing of the fusion protein are shown as boxes in blue and red (if different) or in green (if the fusion comprises a repeat of the same monomer). The two purple domains at the N-terminus are transmembrane domains (TMDs) as predicted by TMHMM, the first TMD is a part of the signal peptide, the second domain is the anchoring TMD that follows the chloroplast transit peptide (Class I targeting presequence). Contig IDs from the presented transcriptome are given for *E. longa* models (EL), accessions are listed for their *E. gracilis* orthologs (EG); GEFR and GDJR accessions are from TSA records GEFR00000000.1 [7] and GDJR00000000.1 [9], respectively.

Moreover, we found four *E. longa* plastid proteins that apparently reach the organelle as translational fusions with other proteins endowed with an N-terminal targeting presequence. Specifically, we observed ribosomal proteins L10 and L17 fused to the C-terminus of ribosomal proteins L15 and L28, respectively, methionyl-tRNA synthetase fused to the C-terminus of tyrosyl-tRNA synthetase, and a putative dicarboxylate carrier fused to the C-terminus of superoxide dismutase (Fig. 3). All of these fusion proteins have a similar structure, comprised of the N-terminal BTS followed by protein monomers separated by a short peptide linker. Hence, unlike the RBCS multimer, these transcripts encode two different proteins. All these fusions are encountered also in *E. gracilis* (Fig. 3) and are thus unlikely to be assembly artefacts. The list of proteins delivered to euglenophyte plastids as translational fusions will probably grow with a more in-depth analysis. For example, while investigating the family of FTSH proteases (see the next section), we found out that one of the predicted plastid-targeted paralogs shared by *E. gracilis* and both *Eutreptiella* species (yet absent from *E. longa*) has a C-terminal extension corresponding to the uncharacterized plastid protein Ycf45 (in some algae encoded by the plastid genome) (Fig. 3). Whether the fused proteins are processed in the plastid by cleavage or remain joined together needs to be determined.

### Euglenophyte and *E. longa*-specific simplifications of the basic plastid infrastructure

The fact that euglenophyte presequences include a region with characteristics of the plastid transit peptide implies the existence of a plastid import machinery homologous to the translocon of the outer/inner chloroplast membrane (TOC/TIC) of other plastids [14]. However, we failed to identify homologs of most of the TOC/TIC components in the *E. longa* transcriptome even when using HMMER and profile HMMs for the respective protein families (i.e. an approach substantially more sensitive than conventional BLAST). The only exceptions were the proteins TIC32 and TIC62, which belong to a large family of short-chain dehydrogenases [48, 49]. TIC32 was described as a calmodulin-binding, NADPH-dependent regulator of the plant TIC, operating in a redox- and calcium-dependent manner [49]. Proteins (with a putative plastid BTS) highly similar to the plant TIC32 are found in *E. longa* as well as other euglenophytes (Fig. 4; Additional file 2: Table S3), and a phylogenetic analysis places them closer to the plant TIC32 than to other related proteins (data not shown), suggesting their functional equivalence. However, little is known about TIC32 and its TIC-independent function is conceivable. In contrast, even the most similar euglenophyte homologs of the plant TIC62 do not cluster with them in a phylogenetic analysis (data not shown), indicating that they should not be considered as candidates for TIC components.

Surprisingly, the transcriptomes of *E. gracilis* and *Eutreptiella* spp. proved to encode discernible homologs of only two additional plastid translocon subunits, TIC21 and TIC55 (Fig. 4; Additional file 2: Table S3). TIC21 (three copies in *E. gracilis* and one in *Eutreptiella* spp.) is only loosely associated with the central translocon subunits and is employed mainly for the import of photosynthesis-related proteins, whereas the import of several non-photosynthetic housekeeping proteins was shown to be unimpaired in TIC21-depleted plant plastids [50]. TIC55 was recently shown to serve as phyllobilin hydroxylase in the chlorophyll breakdown pathway and its role in plastid protein import was questioned [51]. Regardless, neither of the euglenophyte TIC55-related proteins is a *bona fide* TIC55 ortholog, as demonstrated by our phylogenetic analysis (Additional file 1: Figure S5). Three sequences correspond to chlorophyllide *a* oxygenase (CAO), an enzyme of chlorophyll *b* synthesis, whereas the others represent different branches within a broader radiation of plant, algal and cyanobacterial proteins including not only TIC55, but also PAO (pheophorbide *a* oxygenase) and PTC52 (a potential chlorophyllide *a* oxygenase).

**Fig. 4:**
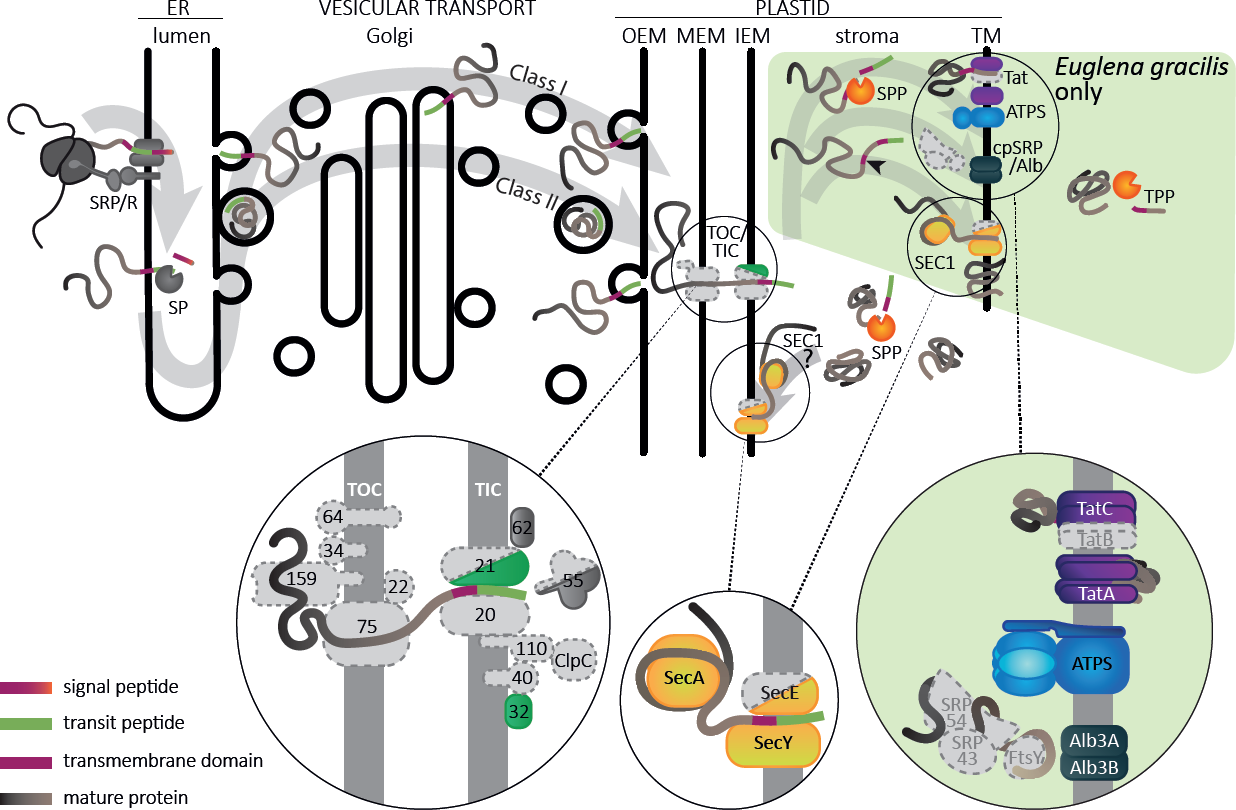
A hypothetical scheme of translocation of plastidial proteins in *Euglena*. The route of plastidial proteins in the *Euglena* cell is schematically depicted, with a zoom-in on individual translocon complexes and protein subunits found in the transcriptomic data analysed. Note that transport across the thylakoid membrane presumably occurs only in the photosynthetic *E. gracilis*, as it is unknown whether the *E. longa* plastid has thylakoids, too. This transport path is taken by proteins with an extended BTS including a third transmembrane domain-like region (arrowhead). The colour code for peptide subdomains is shown in the lower left corner. Note that receptor GTPases Toc34 and Toc159 represent in the figure broader families of paralogous proteins including Toc33 and Toc120/Toc132, respectively. OEM/MEM/IEM/TM: plastid outer, middle, inner envelope membranes and thylakoid membrane; SRP/R: signal recognition particle (receptor) complex; SP: signal peptidase; TOC/TIC: translocon of the outer/inner chloroplast membrane; SPP/TPP: stromal/thylakoid processing peptidase; ATPS: ATP synthase.

While the apparent absence of TIC21 and TIC55-related proteins in *E. longa* is obviously related to the loss of photosynthesis, the lack of discernible homologs of core TOC/TIC components in euglenophytes in general is striking. It is possible that euglenophytes still possess a form of the TOC/TIC translocon, yet with its components diverged beyond recognition by the bioinformatics tools employed by us. Another possibility is that the original protein import machinery of the green algal progenitor of the euglenophyte plastid was replaced by a novel apparatus that acquired the ability to sort proteins according to similar characteristics (i.e. the presence of an N-terminal plastid transit peptide) as the conventional TOC/TIC translocon. This would not be without a precedent. In algae with rhodophyte-derived four membrane-bound plastids, the N-terminal transit peptide-like region not only enables import into the plastid stroma via the TOC/TIC translocon, but first serves as a sorting signal (recognized by a hitherto uncharacterized receptor) for translocation of the preprotein across the second outermost plastid membrane into the periplastid space mediated by a unique machinery called SELMA [52].

After a protein has passed into the plastid, its presequence needs to be cleaved off by the stromal processing peptidase, conserved in *E. longa* as well as other euglenophytes (Additional file 2: Table S3). Photosynthetic plastids need to translocate specific proteins further into the lumen or the membrane of thylakoids. Several different machineries mediating this step are known, including the Tat (twin-arginine translocase), SRP/Alb3 (Signal Recognition Particle/Albino3) system, and the SEC translocase [53]. Critical subunits of the Tat translocase (TatA and TatC) and two different forms of the Alb3 protein can be readily identified in the transcriptome of *E. gracilis* and *Eutreptiella* spp., whereas no such homologs can be found in the *E. longa* transcriptome assembly (Fig. 4; Additional file 2: Table S3). This is consistent with the role of these proteins in translocating exclusively the components of the photosynthetic machinery. In addition, the Tat translocase depends on the electric potential across the thylakoid membrane [54], and hence would be useless in plastids lacking a mechanism to generate it. In non-photosynthetic plastids the transmembrane electrochemical proton gradient is maintained by the ATP synthase at the expense of ATP. Indeed, Tat is missing from non-photosynthetic plastids that lost ATP synthase [31]. The absence of both Tat and ATP synthase in *E. longa* is thus fully consistent with these insights. More unexpected is the absence of the chloroplast signal recognition particle (cpSRP) components (cpSRP54 and cpSRP43) and its receptor (cpFtsY) in all euglenophytes (Fig. 4), since these proteins are conserved in photosynthetic plastids in general (cpSRP54 and cpFtsY) or in plants and green algae (cpSRP43) [55, 56] and are important for the delivery of substrate proteins to the Alb3 insertase [53]. How the absence of cpSRP/cpFtsY affects the function of the euglenophyte Alb3 remains to be elucidated.

In contrast, the third plastid translocase, SEC, is conserved in both *Euglena* species as well as in *Eutreptiella* spp. (Fig. 4; Additional file 2: Table S3). Two different plastid SEC systems were described in plants, one located in thylakoid membranes and the other in the chloroplast inner envelope membrane (IEM) [57]. We retrieved only a single SecA and SecY subunit homolog in each *Euglena* species, and in *E. gracilis* we found a single homolog of the third SEC subunit, SecE, whereas *E. longa* apparently lacks it (its absence was confirmed by searching raw RNA-seq reads). Our phylogenetic analyses revealed that the euglenophyte SecY and SecA proteins are related to the plant and green algal components of the thylakoid-associated SEC1 system (Fig. 5); the phylogeny of the short and poorly conserved SecE protein was not analysed. The apparent absence of the SEC2 complex in euglenophytes is intriguing, but can be explained by at least three different scenarios: (1) the SEC1 complex is in fact not exclusive for thylakoids and operates also in the IEM to facilitate insertion of some IEM proteins, whereas the SEC2-specific substrates have been lost from euglenophytes; (2) SEC1 operates also at the IEM and has taken over some of the SEC2 substrates; (3) there is no SEC machinery at the IEM.

**Fig. 5:**
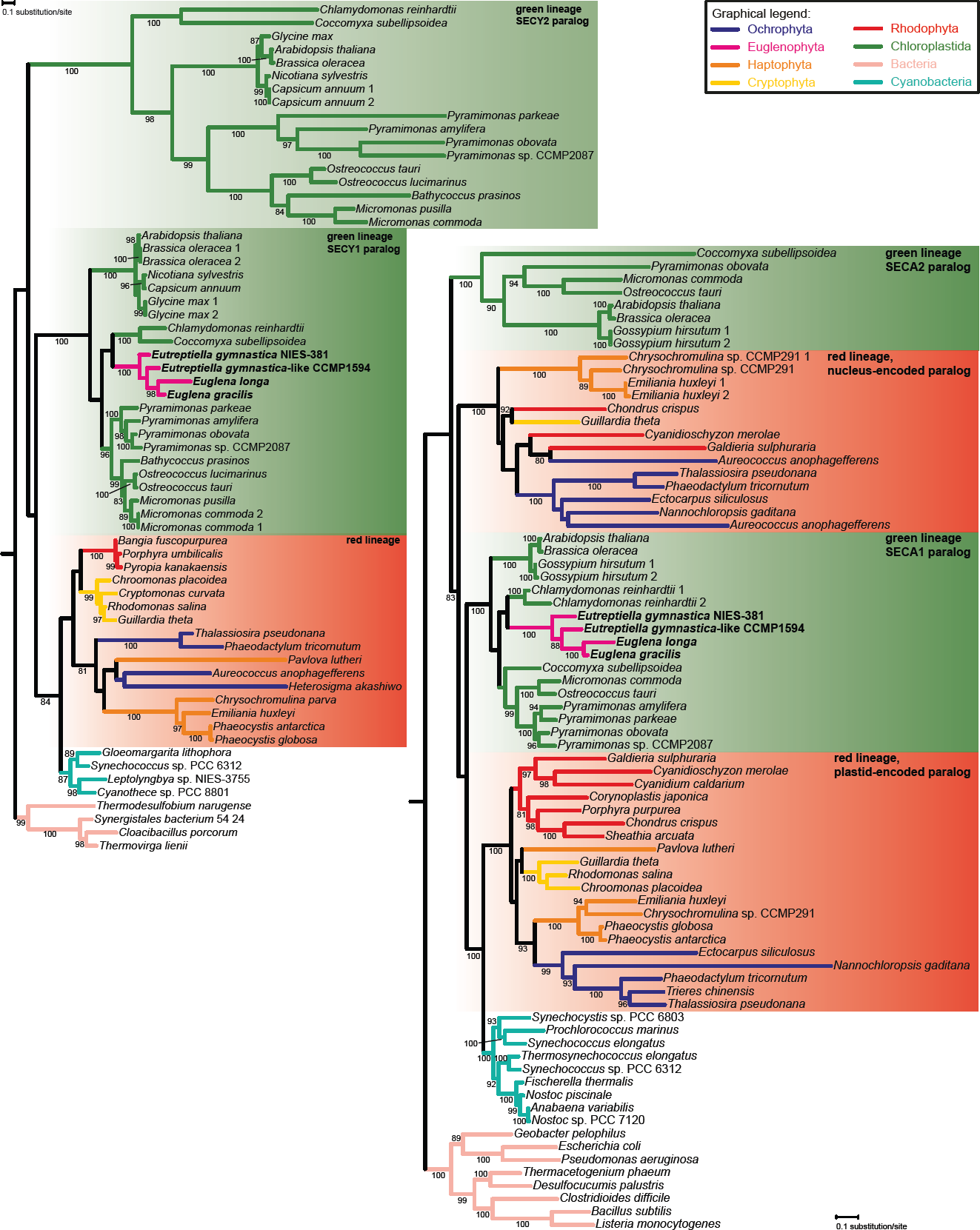
Phylogenetic analysis of the SecA and SecY subunits of the SEC translocon. The maximum likelihood tree of SecA and SecY proteins documents that the euglenophyte proteins are orthologs of the chlorophyte SECA1 and SECY1, respectively. Bootstrap support values are given when ≥ 80.

Studies in plant plastids have so far identified only three putative SEC2 substrates, the TIC complex components TIC40 and TIC110 (both apparently missing from euglenophytes) and the FTSH12 protein, one of the paralogs of an expanded family of membrane-bound proteases [58]. We identified FTSH protease homologs in *E. longa* and the three photosynthetic euglenophytes and performed a phylogenetic analysis by including the well annotated FTSH protease set from *Arabidopsis thaliana* (Additional file 1: Figure S6). Like their *A. thaliana* homologs [59], the euglenophyte FTSH proteases are presumably mitochondrion- or plastid-targeted and their predicted localisation is highly congruent with the phylogenetic relationships of the proteins. *E. longa* and *E. gracilis* proved to encode essentially the same set of mitochondrial FTSH proteases (a difference being the existence of two highly similar variants of one homolog in *E. longa* and both *Eutreptiella* species, possibly due to very recent gene duplication). In comparison, *E. longa* possesses a reduced complement of plastid-localised FTSH proteases compared with the photosynthetic euglenophytes (three versus six to nine), most likely due to the loss of paralogs specialized to act on photosynthesis-related proteins. Interestingly, the plastid-localised FTSH proteases of euglenophytes all group with *A. thaliana* homologs from the thylakoid membrane (FTSH 1, 2, 5, and 8) [59], hence our *in silico* analysis does not recover any obvious FTSH protease candidates to localise to the euglenophyte IEM. Crucially, the absence of a euglenophyte ortholog of FTSH12 (Additional file 1: Figure S6) is consistent with the lack of the SEC2 complex.

Nevertheless, it is still possible that some of the hitherto unidentified SEC2 substrates have been preserved in euglenophytes, but their import was taken over by SEC1. Operation of the same SEC translocon in both the thylakoids and the IEM was the primitive state in plastid evolution as documented by the arrangement in cyanobacteria [60] and glaucophytes [61]. Furthermore, the presence of the SEC1 complex in *E. longa* also supports its localisation to the IEM, since this non-photosynthetic species is unlikely to have thylakoids (although this needs to be proven by electron microscopy). The apparent lack of a plastidial SecE homolog in *E. longa* may reflect functional simplification of the translocase associated with the loss of its predominant substrates (i.e. proteins of the photosynthetic machinery). Finally, it is possible that no SEC machinery is located in the IEM of the euglenophyte plastids and all proteins residing in this membrane (presumably a number of metabolite transporters and components of the elusive protein import machinery) reach their destination via the so-called stop-transfer pathway, i.e. lateral insertion into the IEM during import of the protein [53]. Whereas in most studied plastids this pathway is utilized by only a subset of IEM proteins, the apparently unusual protein import apparatus in euglenophyte plastids (see above) might suggest that this mechanism serves as a general route for the IEM proteins delivery.

We also used our transcriptome assembly to investigate whether *E. longa* has preserved the conventional plastid division machinery, comprising proteins functioning outside (e.g. the dynamin family GTPase Arc5/DRP5B) and inside (e.g. the tubulin homolog FtsZ and Min proteins) the plastid [62]. None of these proteins could be identified in our data, and *E. gracilis* and *Eutreptiella* spp. also appear to lack them (Additional file 2: Table S3). The absence of Arc5/DRP5B is not extraordinary, since this protein is also missing in the sequenced representatives of glaucophytes, cryptophytes, chlorarachniophytes, and myzozoans [63]. The absence of FtsZ in *E. longa* would not be particularly surprising either, since myzozoans with non-photosynthetic plastids (apicomplexans and *Perkinsus marinus*) are devoid of it, too. However, the apparent lack of FtsZ in euglenophytes in general is noteworthy, because all organisms with photosynthetic plastids studied to date do keep FtsZ, typically as multiple paralogs [63]. Our analyses thus suggest that the original plastid division mechanism of the green algal donor was substantially simplified or modified during the endosymbiotic integration of the euglenophyte secondary plastid.

### A plastid-targeted Rho factor homolog in euglenophytes acquired by HGT from bacteria

The plastid genomes of euglenophytes including *E. longa* encode four subunits of the RNA polymerase responsible for its transcription, namely alpha (RpoA), beta (RpoB), beta’ (RpoC1), and beta’’ (RpoC2) (Additional file 1: Figure S3). The fifth putative subunit, i.e. the sigma factor (RpoD), is found in the transcriptome of *E. longa*, *E. gracilis*, and both *Eutreptiella* species (Additional file 2: Table S3), completing the conventional cyanobacteria-derived RNA polymerase holoenzyme conserved in plastids in general [64]. However, while surveying the *E. longa* transcriptome for potential plastid-targeted proteins we unexpectedly encountered a homolog of the Rho factor, a highly conserved and widespread component of eubacterial transcription machinery [65] that, to the best of our knowledge, has never been reported from eukaryotes. This is evidently not due to a bacterial contamination. The respective contig carries the characteristic SL sequence at its 5’-end (Additional file 2: Table S3), the encoded protein has an N-terminal, BTS-like extension compared to the bacterial homologs (Additional file 1: Figure S7), and closely related homologs exist in *E. gracilis* and both *Eutreptiella* species (although the *E. gymnastica*-like CCMP1594 sequence is truncated and the 5’-end could not be completed by iterative read mapping) (Fig. 6; Additional file 2: Table S3).

**Fig. 6:**
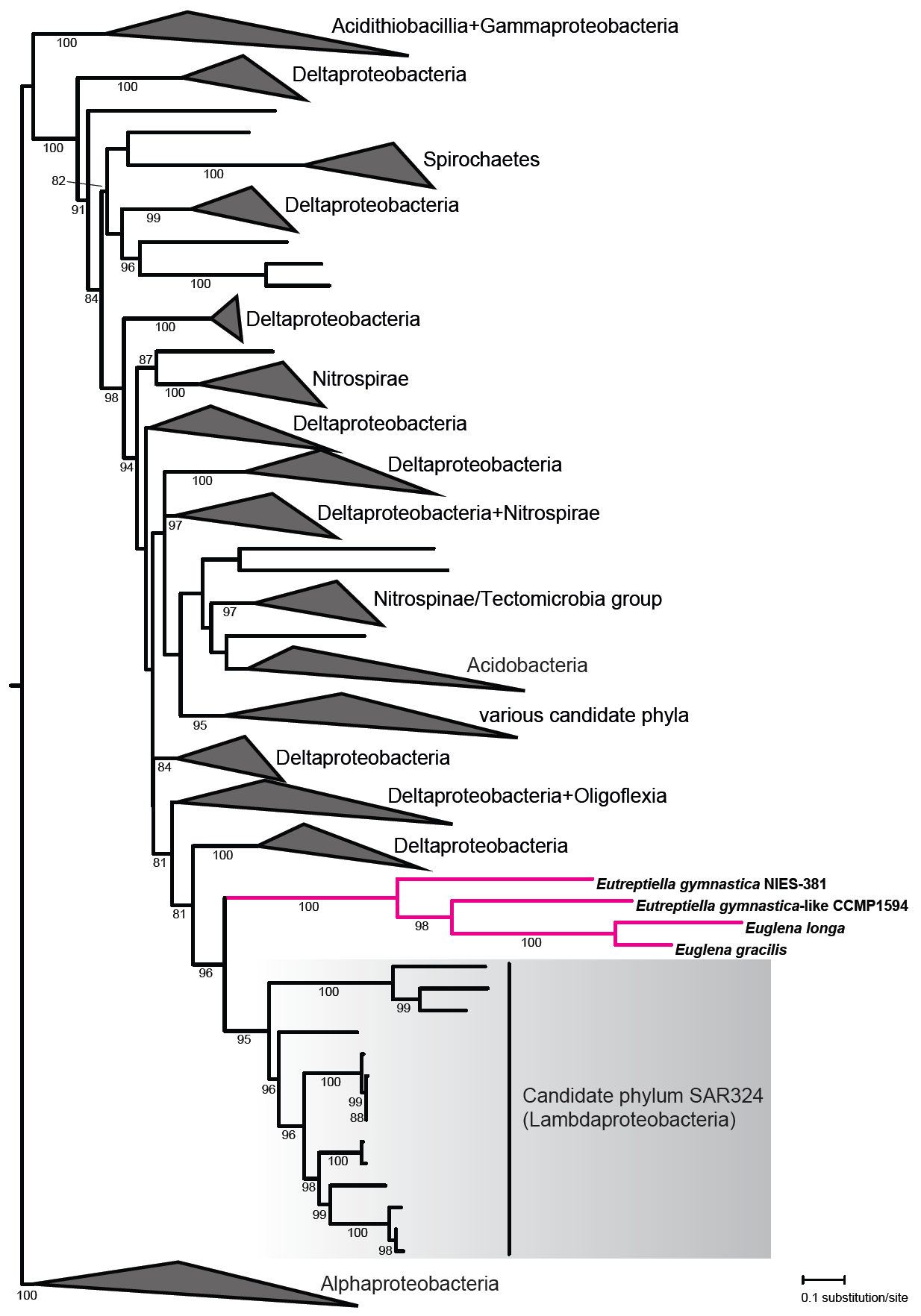
The phylogenetic position of the euglenophyte translation-termination factor Rho among bacterial homologs. The tree was inferred by using the maximum likelihood method. Bootstrap support values are given when ≥ 80. For simplicity, various broader clades were collapsed and the taxonomic provenance of the sequences included in them is indicated at the triangles. A full version of the tree is provided in the Additional file 3).

The Rho factor is a homohexameric ATP-driven RNA helicase that is critical for proper transcription termination of a sizeable proportion of genes in bacteria [65]. Despite being so important and prevalent, it has not been retained in the endosymbiotic organelles of eukaryotes; the few eukaryotic hits retrieved by a blastp search against the NCBI nr protein database are all contaminants from bacteria (Additional file 2: Table S6). In the case of the plastid this seems to be due to an earlier Rho factor loss in Cyanobacteria, as documented by our searches that failed to identify convincing Rho factor homologs in this bacterial phylum (the few hits all seem to be contaminants from other bacteria; Additional file 2: Table S6). Hence, the emergence of the Rho factor in the euglenophyte plastid is indeed striking. Our phylogenetic analysis indicates that the donor of the euglenophyte Rho factor was related to the recently recognized bacterial phylum SAR324 ([66]; see also http://gtdb.ecogenomic.org/) (Fig. 6; Additional file 3). This bacterial lineage (also called Candidatus Lambdaproteobacteria) so far lacks cultured representatives and all available genomic data are derived from metagenomes or single-cell genome sequencing (e.g. [66–70]). The euglenophyte Rho factor thus represents an interesting example of a HGT-derived eukaryotic gene whose actual bacterial source could be properly identified only owing to the recent substantial improvement of the genome sampling of the bacterial phylogenetic diversity.

What might be the function of the Rho factor in the euglenophyte plastid? Sequence comparison of the euglenophyte and bacterial Rho proteins reveals that the motifs critical for their function have not been affected by the gene transfer into the eukaryotic lineage (Additional file 1: Figure S7). Therefore, there is no reason to assume that the function of the euglenophyte Rho factor is different from the bacterial prototype at the biochemical level and the obvious null hypothesis is that the euglenophyte Rho factor is involved in transcription termination in the plastid. The Rho factor in bacteria terminates transcription in only a specific subset of genes, but a common Rho-specific termination signal was not found [65]. Hence, it cannot be decided at the moment which euglenophyte plastid genes are the best candidates for being regulated by the Rho factor. Future biochemical experiments should enable us to test whether the Rho factor is involved in transcription termination in euglenophyte plastids and if so, which genes are its specific targets.

## Conclusionss

We have sequenced and assembled the transcriptome of an interesting organism and a useful model for studying plastid reduction accompanying the loss of photosynthesis – a widespread phenomenon among non-photosynthetic plants, algae and protists [71]. Our analyses suggest that our *E. longa* transcriptome assembly provides a good representation of nucleus-encoded plastid-targeted proteins in general. We also confirmed that the N-terminal plastid-targeting presequences in *E. longa* exhibit the same characteristic structure as in *E. gracilis*, which opens up a possibility of a systematic bioinformatic survey of the *E. longa* plastid proteome. Complete reconstruction of the metabolic pathways localized in the non-photosynthetic plastid of *E. longa* will help to understand its physiological role(s). Work on this task is in progress in our laboratories.

In this paper, we exploited the transcriptome assemblies of *E. longa* and its photosynthetic relatives to illuminate the molecular machinery responsible for plastid biogenesis and division. As expected, we observed selective loss of components linked to the loss of photosynthesis in *E. longa*, but strikingly, our results revealed that euglenophytes as a whole have lost many components present in the green algal donors of their plastid and conserved in plants and algae in general. Most notable is the elusive nature of the euglenophyte mechanisms of plastid protein import and plastid division. On the other hand, the identification of the plastid-targeted Rho factor acquired by HGT from bacteria points to an enrichment of the molecular machinery of the euglenophyte plastid unprecedented among eukaryotes. Previous phylogenetic analyses unveiled a mosaic nature of the euglenophyte plastid proteome, indicating that many proteins, such as some enzymes of the Calvin cycle or the MEP pathway of isoprenoid biosynthesis, were gained by horizontal gene transfer (HGT) from various algal sources different from the donor of the plastid itself [17, 18, 72]. Our results suggest that the euglenophyte plastid proteome has an even more complex evolutionary origin, including a contribution from bacteria. Our results thus emphasize the need to revive the interest in how the euglenophyte plastids have evolved and function as cellular organelles.

## Materials and Methods

### Culture conditions, RNA isolation and mRNA extraction, cDNA synthesis, and PCR

*Euglena longa* strain CCAP 1204-17a was cultivated statically in the dark or under constant illumination at 26 °C in Cramer-Myers medium [73] supplemented with ethanol (0.8% v/v). The cultures were not completely axenic, but the contaminating bacteria were kept at as low level as possible. RNA was isolated using TRIzol^®^ Reagent (Invitrogen, Carlsbad, USA) and mRNA was then extracted using PolyATtract mRNA Isolation Systems III (Promega, Madison, USA). cDNA synthesis was carried out with an oligo(dT) primer using Transcriptor First Strand cDNA Synthesis Kit (Roche, Basel, Switzerland). Sequences of all primers used in PCR experiments are listed in Additional file 2: Table S1. PCR products were amplified from 30 ng of *E. longa* cDNA using MyTaq™ Red DNA Polymerase (Bioline, London, UK). The presence of SL-sequence at the 5’-end of transcripts was tested using the same PCR experimental design as described previously [28, 74]. PCR conditions were as follows: 95°C for 1 min; 35 cycles of 95°C for 15 sec, 50°C for 15 sec, 72°C for 1 min, and the final extension at 72°C for 5 min. The PCR products were purified (Gel/PCR DNA Fragments Extraction Kit, Geneaid Biotech, New Taipei City, Taiwan) and their identity was verified by sequencing (Macrogen Europe, Amsterdam, Netherlands).

### *E. longa* transcriptome sequencing and assembly

Library preparation and sequencing of dark-grown and illuminated *E. longa* was performed by GATC Biotech on an Illumina HiSeq 2000 platform. A total of 47,442,811 paired-end 80-bp reads were obtained. Contamination from *Homo sapiens* and *Capsicum annuum* identified in a preliminary transcriptome assembly was removed by mapping the reads to the genome sequences of the respective species using Deconseq 0.43 [75]. The remaining reads were adapter- and quality-trimmed by Trimmomatic 0.33 [76]. The final read assembly was performed using the ABySS software 1.52 (k-mers 31-51; [77]), then fused with Trans-ABySS 1.48 [78], Trinity r20140717 (k-mers 31 and 25; [79]), and SOAPdenovo-Trans 1.04 (k-mers 31 and 33; [80]), followed by merging the contigs (>99% sequence identity over 150 nt) using CAP3 12/21/07 [81]. The completeness of the assembly was assessed by a BUSCO search of conserved eukaryotic orthologs using the transcript mode and eukaryotaV1 and eukaryotaV2 sets of orthologs [82].

### Sequence searches and phylogenetic analyses

Homologs of proteins of interest were searched in the final transcriptome assembly using local tBLASTn [83]. The contigs representing candidate hits were translated in all six frames and the corresponding protein model was selected. To possibly detect sequences not identified by tBLASTn, we employed HMMER 3.1b2, a more sensitive method of homology detection based on profile hidden Markov models [84]. Profile HMMs were built from seed alignments of the proteins families of interest obtained from the Pfam database and used to search the transcriptome of *E. longa* and both available *E. gracilis* transcriptomes (accession numbers GDJR00000000.1 and GEFR00000000.1; [9] and [7], respectively) translated in all six frames by an in-house python script, and of both *Eutreptiella* species (reassemblies available at zenodo.org DOI: 10.5281/zenodo.257410). Searches for candidates for TOC/TIC machinery components in euglenophytes were in parallel done by iterative HMMER searches to enable identification of even more distant homologs. Specifically, alignments of full proteins from the RefSeq database and alignments of separate domains from the Conserved Domains Database (CDD; [85]) were used to construct the initial profile HMMs for searching transcriptomes of several chlorophytes including *Pyramimonas parkeae* and *Pyramimonas obovata* (sequenced in frame of the MMETSP project [10]), which represent close relatives of the putative euglenophyte plastid donor [1, 11–13]. The identified chlorophyte homologs were re-aligned with sequences of the initial reference set using ClustalW [86], new profile HMMs were built and euglenophyte sequence data were searched with them. To further confirm the absence of some genes in the transcriptome of *E. longa*, tBLASTn searches were carried out against the unassembled raw reads using the respective protein sequences from *E. gracilis* as queries. Iterative searches of raw reads were also used in attempts to extend termini of contigs that proved to include truncated coding sequences (taking into account also linking information provided by pair-end reads).

Based on the analysis of high-confidence candidates for *E. longa* plastid-targeted proteins and the known structure of plastidial BTSs in *E. gracilis*, the criteria for identifying a protein as plastid-targeted were set as follows: (1) the signal peptide was predicted by the PrediSi [87] or PredSL [88] programs; (2) one or two transmembrane domains at the N-terminus of the protein were predicted by the TMHMM program [89] available online or implemented in Geneious 10.1.3 [90]. The resulting set of sequences was further filtered by checking for the presence of a plastid transit peptide, which was predicted by MultiLoc2 [91] after *in silico* removal of the signal peptide or the first transmembrane domain.

Phylogenetic analyses were carried out for selected proteins. Homologs were identified by BLAST searches in the non-redundant protein sequence database at NCBI and protein models of selected organisms from JGI (Joint Genome Institute, jgi.doe.gov), Ensembl (www.ensembl.org), and MMETSP (Marine Microbial Eukaryote Transcriptome Sequencing Project; original assemblies from marinemicroeukaryotes.org and reassemblies currently available at zenodo.org DOI: 10.5281/zenodo.257410; [10]). Sequences were aligned using the MAFFT 7 tool [92] and poorly aligned positions were eliminated with the trimAL tool [93]. The alignments were manually refined using AliView [94] and ambiguously aligned positions were removed. For presentation purposes alignments were processed using the programme CHROMA [95]. Maximum likelihood (ML) trees were inferred from the alignments using the best-fitting substitution model as determined by the IQ-TREE software [96] and employing the strategy of rapid bootstrapping followed by a “thorough” ML search with 1,000 bootstrap replicates. The list of species, and the number of sequences and amino acid positions are present in Additional file 2: Tables S7-12 for each phylogenetic tree.

## Abbreviations

BTS: bipartite topogenic signal
HGT: horizontal gene transfer
IEM: inner envelope membrane
SL: spliced leader
TMD: transmembrane domain

## Declarations

### Ethics approval and consent to participate

Not applicable.

### Consent for publication

Not applicable.

### Availability of data and materials

The raw sequencing data and the final assembly of the *E. longa* transcriptome are available at NCBI (www.ncbi.nlm.nih.gov) as BioProject PRJNA471257. The multiple sequence alignments used for phylogenetic analyses are available upon request from the corresponding author.

### Authors’ contributions

KZ and ZF performed most bioinformatic analyses and prepared figures and tables. AMGNV contributed by analyses of the TOC/TIC system. KZ, MV, and JK maintained the *E. longa* culture and prepared RNA for transcriptome sequencing. EB and VK processed and assembled the RNAseq data. MO contributed to the design of the study and the paper. ME conceived the study, performed some bioinformatics analyses and prepared the first draft of the manuscript. All authors edited the manuscript and approved its final form.

## Acknowledgements

We thank Lieven Sterck (Department of Plant Biotechnology and Bioinformatics, Ghent University) for his help with obtaining RNA-seq data from *E. longa*. We acknowledge computation resources provided by CERIT-SC and MetaCentrum, Brno, Czech Republic.

## Additional files

**Additional file 1:** Supplementary figures.

**Additional file 2:** Supplementary tables.

**Additional file 3:** Full tree of the termination factor Rho in the Newick format.

